# A high throughput screen for active human transposable elements

**DOI:** 10.1101/192708

**Authors:** Erika M. Kvikstad, Paolo Piazza, Jenny C. Taylor, Gerton Lunter

**Affiliations:** Wellcome Trust Centre for Human Genetics, Oxford, UK; National Institute for Health Research Comprehensive Biomedical Research Centre, Oxford, UK; Department of Medicine, Imperial College London, London, UK

**Keywords:** Transposable Elements, *Alu*, LINE1, polymorphism, next generation sequencing, bioinformatics

## Abstract

Transposable elements (TEs) are mobile genetic sequences that randomly propagate within their host’s genome. This mobility has the potential to affect gene transcription and cause disease. However, TEs are technically challenging to identify, which complicates efforts to assess the impact of TE insertions on disease. Here we present a targeted sequencing protocol and computational pipeline to identify polymorphic and novel TE insertions using next-generation sequencing: TE-NGS. The method simultaneously targets the three subfamilies that are responsible for the majority of recent TE activity (L1HS, *Alu*Ya5/8, and *Alu*Yb8/9) thereby obviating the need for multiple experiments and reducing the amount of input material required. Here we describe the laboratory protocol and detection algorithm, and a benchmark experiment for the reference genome NA12878. We demonstrate a substantial enrichment for on-target fragments, and high sensitivity and precision to both reference and NA12878-specific insertions. We report 17 previously unreported loci for this individual which are supported by orthogonal long-read evidence, and we identify 1,471 polymorphic and novel TEs in 12 additional samples that were previously undocumented in databases of insertion polymorphisms. We anticipate that future applications of TE-NGS alongside exome sequencing of patients with sporadic disease will reduce the number of unresolved cases, and improve estimates of the contribution of TEs to human genetic disease.

## INTRODUCTION

Genome sequencing is now routinely used to identify the mutations responsible for rare genetic disease. Recent large-scale sequencing efforts of individuals with rare Mendelian disorders indicate that in approximately 20-40% of cases a causal variant can be identified (eg, [1-4]). To date, these efforts have focused mainly on small variants in exonic regions. This leaves open the possibility that a substantial fraction of the remaining causal variants are localized in non-coding regions and affect gene regulation, rather than change the proteins themselves.

One class of mutations with a potentially large effect on gene regulation are transposable element (TE) insertions. TEs are self-replicating mobile elements that randomly insert new copies of themselves into their host’s genome, with the result that in modern humans up to 60% of the genome ultimately derives from TE insertions [5, 6]. Most of these insertions were “dead on arrival” or have been inactivated over time, and today only a small collection of loci are thought to be active [7-9]. Two of these active classes of retrotransposons, LINE1 and *Alu*, copy their sequences to new locations via RNA intermediates [10, 11]. Although the insertion rate of novel TEs is low (∼1 per 20 and 1 per 100 births for *Alu* and LINE1 respectively [12, 13]), together, LINE1 and *Alu* account for 95% of active TE insertion in human genomes [14].

Several recent studies have highlighted the importance of this TE activity for creating population-level sequence diversity [14, 15]. TE-mediated mutations can also cause disease by disrupting genes or modifying their expression. For instance, insertions can directly interrupt exons, causing malformed or partial protein products. In addition, because TEs contain regulatory elements including promoters and transcription start and stop signals, their insertion can cause nearby genes to become dysregulated or truncated (reviewed in [16]).

Because of the relative rarity of TEs and the technical difficulty of identifying them, their contribution to human disease remains unclear. Published estimates suggest that TEs are responsible for a small fraction of human disease (0.27%, [17]), and so far, 96 individual monogenic disease-associated TE insertions have been identified [18]. However, these numbers likely represent an underestimate of the true impact of TEs, in part because they predate the clinical application of next generation sequencing (NGS) technology. Even today, most next generation sequencing (NGS) assays in clinical use involve exome sequencing rather than whole-genome sequencing, and TE insertions in introns and intergenic regions with possible regulatory impact will therefore be missed.

Recently, several targeted NGS technologies have been developed to detect structural variation due to TE insertions [19-25]. Strategies to target TE sequences utilize hybridization approaches in order to selectively enrich for molecules spanning a TE of interest [19, 22, 24]. Alternatively, multiple degenerate primers can be used to amplify unknown flanking sequences in conjunction with a TE-specific primer [21, 25]. Restriction enzyme digestion followed by PCR has also been employed to isolate TE sequences [20, 23]. These strategies require post-hybridization or post-amplification library preparation for NGS sequencing, which substantially increases the amount of input material required (up to 10 μg; [23]); this is important because the amount of available genomic DNA from clinical samples is often limited. In addition, current targeted sequencing approaches typically isolate activities of either *Alu* [22], or LINE1 [20, 21, 23, 25], thus separate methods are required for interrogation of multiple retrotransposons in any one individual [24].

Here we present a sequencing-based whole-genome screen for polymorphic and novel TE insertions that is sensitive, specific, and cost-effective. TE-NGS simultaneously targets three major active TE subfamilies in humans: Human-specific LINE1 (L1HS), *Alu*Ya5/8, and *Alu*Yb8/9, together responsible for the majority (60%) of novel TE insertions [14] (reviewed in [26]). The method creates TE-enriched libraries from genomic DNA or pre-existing genomic DNA libraries, and thus complements whole exome sequencing strategies. The procedure selectively amplifies TE sequence and the flanking regions of actively transposing TE-subfamilies. The protocol includes steps to remove amplified genomic background and sequence artefacts so that very little sequencing capacity is required. A computational pipeline then identifies the signatures of the locations of candidate novel and polymorphic TE loci.

## RESULTS

### Targeted NGS of TE insertions

Figure 1 outlines the main workflow of the TE-NGS protocol (Materials and Methods; see Supplemental Material for detailed procedures). These steps are designed to address several challenges to targeted TE sequencing. First, the combined actively mobile elements constitute a very small fraction of the genome (0.12%, Table S1). Second, active TE of interest are similar to the large fraction of TE-derived genomic sequence [5], and in particular to the relatively abundant but inactive *Alu* and LINE subfamilies *Alu*Sx1 and L1PA2. Third, new *Alu* and LINE1 elements do not insert randomly in the genome [27, 28], resulting in a subset of TE targets (∼0.07%) that are clustered in close proximity and oriented in head-to-head fashion (i.e. inverted repeats; Materials and Methods; [27]), causing issues in the PCR steps of the protocol.

**Fig.1.**
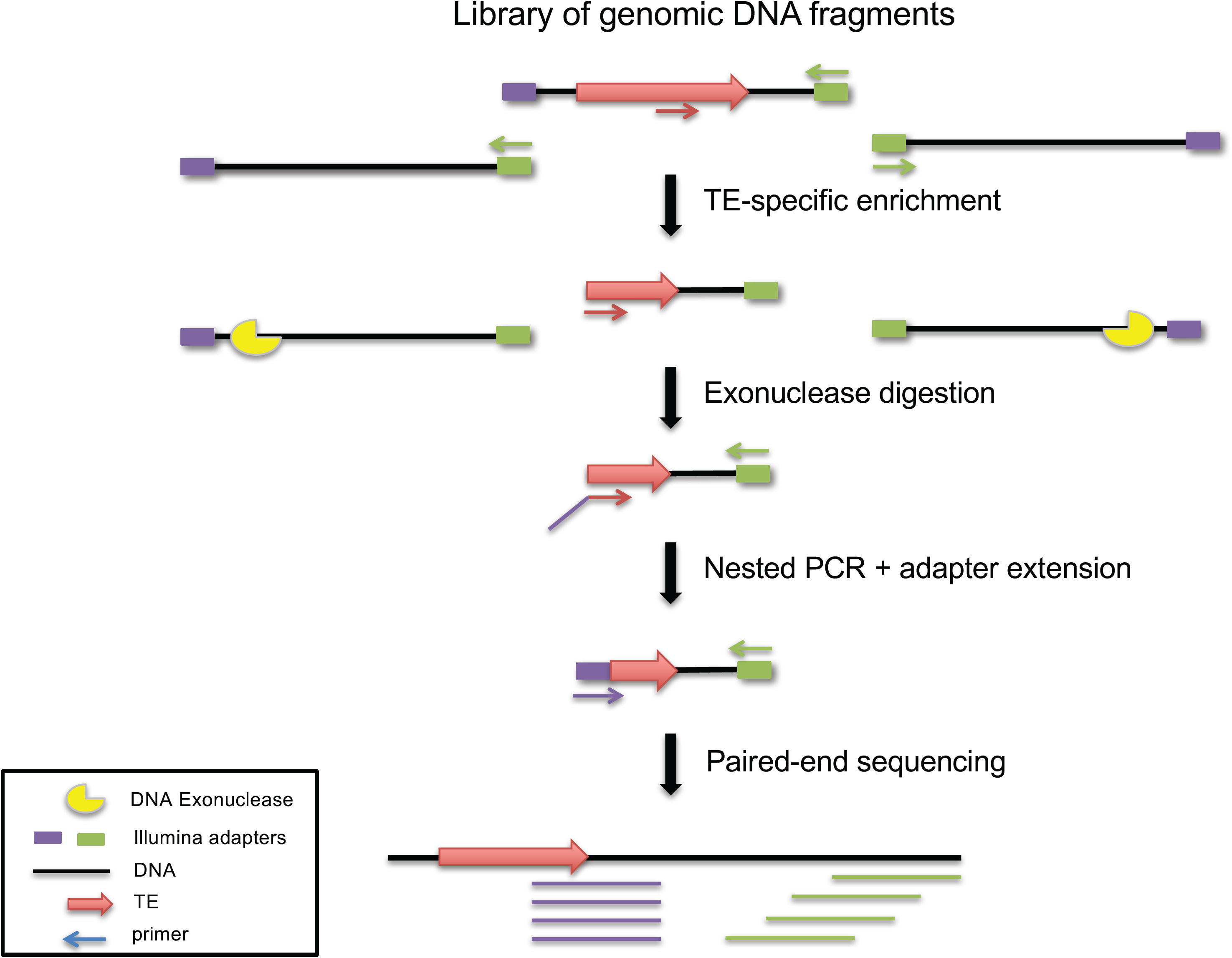
TE-NGS sequencing workflow. Enrichment for genomic fragments spanning active TEs and their unique flanking sequence is achieved by several enzymatic steps as described in the main text. First, genomic DNA is sheared, and adapters for sequencing are ligated to the genomic fragments following standard library preparation protocols. Next, a small aliquot (10 ng) of library is used as template for targeted amplification with primers complementary to TE subfamily-specific sequences and to the Illumina Universal PCR (P5) primer. Remaining genomic background fragments are removed by ssDNA exonuclease digestion after linear amplification with TE-target primers. Finally, amplification with nested primers targeting TE diagnostic bases, and containing Illumina i7 index and P7 primer sequences generates full double-stranded dual-adapter libraries containing unique indices for each sample and each TE subfamily, allowing for downstream pooling and multiplexing of many samples simultaneously (see Materials and Methods; Supplemental Material for detailed procedures).

To overcome these challenges, we first prepare genomic libraries using standard NGS procedures followed by PCR enrichment of library fragments containing TE insertions and their flanking regions. In a single reaction, two TE-target primers are used, each designed to anneal to bases present in the *Alu* (*Alu*Yb8/9 and *Alu*Ya5/8), and L1HS subfamilies, respectively (Table S2). These primers are complementary to the 3’ ends of TE insertions, which contain motifs that distinguish the active subfamilies from closely related but inactive family members. Exponential amplification is achieved by multiplex PCR of the two TE-target primers each in conjunction with the Illumina Universal (P5) primer that will anneal to the complimentary i5 adapter present on all library molecules. Thus, many loci in the genome are amplified simultaneously without requiring degenerate primers.

Targeted sequences are expected to form at most a third of the product after this step, because in addition to exponential amplification of target elements, abundant background genome fragments lacking TE primer sites but containing the Illumina adapters (and P5 priming sites) are linearly amplified at each cycle. Additionally, target elements oriented in a head-to-head fashion will also amplify exponentially, since both ends of the fragment contain TE-primer sites. These fragments lack complementarity to the i7 Illumina sequencing primer necessary for cluster formation via bridge sequencing, and therefore do not yield usable data, but deplete the primer pool and cause issues in library normalization, affecting sequencing.

To remove these unwanted PCR products, TE-NGS libraries are processed by two rounds of asymmetric amplification and single-stranded (ss)DNA exonuclease digestion. We first perform linear amplification with TE-target primers, leaving genomic background molecules lacking the TE largely in single-stranded form. Next, digestion by a ssDNA exonuclease selectively removes these molecules to produce a pool of dsDNA fragments containing TE targets. Similarly, molecules spanning inverted repeats are removed from the library by linear amplification with the Illumina Universal primer, so that fragments containing two TEs in head-to-head orientation, which after amplification lack adapter sequence, remain in a ssDNA configuration. After a second incubation with ssDNA exonuclease, these molecules are removed, leaving the resulting library consisting largely of fragments overlapping a single TE-target locus and its unique flanking sequence.

Finally, specificity is increased by three individual nested PCR reactions, each with a nested primer targeting a specific subfamily. At this step an Illumina index adapter is incorporated, thus producing TE-enriched libraries ready for multiplexing and sequencing by paired-end reads using Illumina platforms (Illumina Inc, San Diego CA; see Fig. S1 for representative library fragment profiles).

### TE identification pipeline

TE detection from targeted NGS sequencing data is performed by a custom bioinformatics pipeline (Fig. S2). First, reads are aligned to the reference genome. Next, they are filtered by requiring the presence of TE-nested primers, in addition to the presence of downstream “TE-like” sequence (see Materials and Methods for details), and further filtered for mapping quality to avoid spurious calls. The remaining reads are then clustered by genomic position. The resulting clusters are annotated using a collection of *Alu* and LINE insertions taken from databases of known reference TEs [29, 30] and of previously published polymorphic germline TE insertions [14, 15, 21, 22, 31]; the latter is referred to as the polymorphic TE database (polyTEdb).

Calls produced by the above pipeline are classified as *reference* if the reference genome contains an insertion that matches the targeted subfamily at that position, and *known non-reference* if a polymorphic TE of the same class (*Alu* or LINE; subfamily-specific information is often not available) at that position is present in polyTEdb. Calls lacking previous evidence of a TE insertion at that position are classified as *novel*. Since a minority of the genome is expected to be inaccessible due to repetitive structures, clusters close to annotated reference gaps (unassembled poly-Ns) or regions enriched for satellite repeats, such as the centromeres and sub-telomeric sequences, and the Y chromosome, were excluded from downstream analysis (Materials and Methods). The detailed procedures, source code and associated annotation files are available on github (https://github.com/ekviky/TE-NGS), with documentation on a github page (https://ekviky.github.io/TE-NGS/).

### Performance evaluation

We applied the TE-NGS workflow to a total of 13 human samples, resulting in 39 TE-enriched libraries (13 samples × 3 TE subfamilies). Individual libraries were pooled and sequenced by 151-bp paired-end sequencing on a single lane of an Illumina Miseq platform (Illumina Inc, San Diego CA, USA), for a total of 25 million reads. On average, we obtained 305,609, 1,343,612, and 337,647 reads per L1HS, *Alu*Ya5/8, and *Alu*Yb8/9 library, respectively (Tables S3, S4). Among the 13 human samples sequenced in this pilot experiment, we included the NA12878 individual originally sequenced by the 1000 Genomes Project and Genome in a Bottle (GiaB) consortia to benchmark the performance of the assay and detection algorithm using previously generated TE call sets [14, 15, 21].

#### Analysis of biochemical assay

We first investigated the ability of the protocol to enrich for active TE families. A substantial fraction of reads from each TE-targeted library contained recognizable matches to the corresponding TE-nested primer (0.922, 0.949 and 0.87 for L1HS, *Alu*Ya5/8, and *Alu*Yb8/9, respectively; Table S3), indicating that genomic background molecules were largely absent from the libraries.

Specificity of the assay to each targeted subfamily was assessed by comparison of observed filtered read coverage of each of the active TEs to closely related subfamilies of high sequence similarity as controls. For example, L1PA2 elements that actively inserted in the common ancestor of human and chimpanzee lineages contain L1HS-nested priming sites that differ by a single nucleotide located at the ultimate 3’ primer position, and are ∼3-fold more abundant in the reference genome than L1HS is (4805 copies, Table S1; [32]). However, we observed 18.68 times more reads mapping to L1HS than L1PA2, and only a small fraction of total L1PA2 loci annotated in the reference genome were covered by 1 or more reads (14%; Table S3). Similarly, *Alu*Sx1, one of the most abundant subfamilies of *Alu*Y (109,589 loci genome-wide; Table S1) was used as control for both *Alu*Ya5/8 and *Alu*Yb8/9 experiments. We observed a substantial enrichment of reads mapping on-target vs. control (19.89 and 594.96-fold for *Alu*Ya5/8 and *Alu*Yb8/9 *vs*. *Alu*Sx1, respectively), and corresponding small proportions of total genome-wide *Alu*Sx1 loci in each *Alu*-enriched library were covered by 1 or more reads (8.7% and 0.5%, for the *Alu*Ya5/8 and *Alu*Yb8/9 libraries respectively; Table S3).

#### Performance of TE detection algorithm

We next evaluated the ability of the detection algorithm to correctly discriminate true TE insertions from false predictions. To do so, we defined two sets of TE insertions to measure the overall sensitivity of the biochemical protocol, and the ability to detect de novo/polymorphic loci in data from a well-characterized reference individual, NA12878. First, we identified a set of 624, 2,739, and 1,847 loci annotated as L1HS, *Alu*Ya5/8, and *Alu*Yb8/9 insertions, respectively, in the GRCh37 reference genome sequence (Reference TEs, Table 1). Since loci located in regions where NA12878 has copy number variants (CNV) could pose alignment ambiguity or be absent altogether, we excluded reference TEs overlapping NA12878-specific CNV calls (Materials and Methods).

**Table 1.**
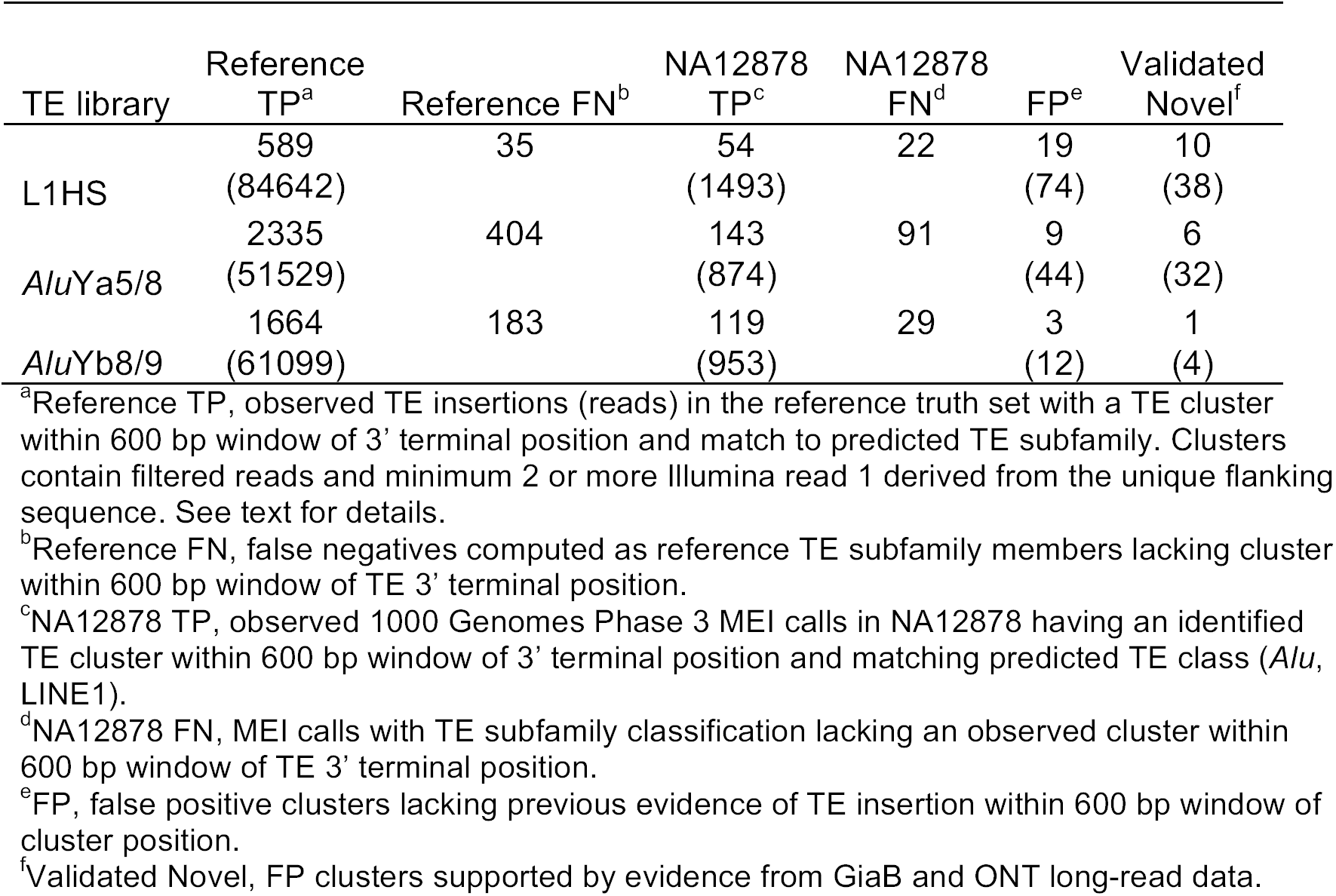
TE loci observed in NA12878 NGS libraries

Second, to assess the sensitivity of the protocol to detect rare and potentially *de novo* loci, we identified a set of polymorphic TE calls. A number of non-reference TE calls for this individual are available through various targeted sequencing detection methods [15, 21], and computational TE discovery methods [14]. However, the limited concordance among methods leaves no clear gold standard set (*e.g.,* Fig. S3, [33]). We therefore obtained 857 polymorphic *Alu* and 76 LINE1 mobile element insertions (“MEI”) calls for NA12878 that were identified by the Phase 3 of 1000 Genomes Project [14], again excluding NA12878-specific regions of CNV. This set of insertions represent the most comprehensive collection of polymorphic TE among the available call sets to date (NA12878 TEs, Table 1).

Given the limited concordance among existing call sets, and the reduced sensitivity of 1000 Genomes calls due to the low fold coverage of sequencing data produced in the project, we assessed specificity of the protocol using the union of the polymorphic NA12878 insertions available. Thus, predictions of our pipeline for which no evidence of a TE insertion at that position exists among all available call sets were considered false positives (FP).

True positives (TP) were defined as predicted calls having a corresponding TE in the set of Reference TEs within 600 bp from the annotated 3’ end of the repeat insertion, and that matches the subfamily annotation. Because only a small number of NA12878 MEI calls in the 1000 Genomes have complete subfamily annotation (163 *Alu*Ya5/8 and 61 *Alu*Yb8/9), we only required a match to the repeat class (*Alu* or LINE1) to consider NA12878 true positives.

False negatives (FN) were defined as Reference TEs and NA12878 TEs annotated as L1HS, *Alu*Ya5/8, or *Alu*Yb8/9 for which no corresponding call in our dataset exists within 600 bp of the annotated 3’ end of the repeat insertion. We restricted 1000 Genomes Phase 3 MEI calls to only those with the *Alu*Yb8/9 and *Alu*Ya5/8 subfamily annotation for determining NA12878 *Alu* false negatives. However, since none of the LINE1 MEI calls have any subfamily designations, in order to be able to compute a NA12878-specific FN rate we conservatively assumed that all 76 of the LINE1 MEI calls were in fact L1HS insertions.

To characterize the ability of TE-NGS to predict true insertions, we evaluated the recall (sensitivity), computed as TP/(TP + FN), for Reference and NA12878 TE sets. Precision, computed as TP/(TP + FP), was used to quantify the ability of the assay to reject false insertion predictions. As illustrated in Figure 2, TE-NGS achieves high rates of recall when considering all loci that are supported by at least one read (0.97, 0.92, 0.94 reference L1HS, *Alu*Ya5/8, and *Alu*Yb8/9, respectively). For example, we observed 33/35 full length MEI LINE1 events annotated in NA12878. However, at this 1-read threshold many spurious calls are made that lack previous evidence of an insertion. We therefore explored precision and recall as a function of cluster read depth (Fig. 2) to determine an effective threshold for defining a TE insertion call. A reasonable balance between sensitivity and specificity was achieved when we required a minimum of 2 reads per cluster derived from the unique flanking sequence 3’ to the insertion site. At this threshold, we observed 0.95, 0.85 and 0.90 Reference (0.71, 0.61, 0.80 NA12878-specific) TE insertions, with precision rates of 0.97, 0.99, and 0.99 for reference calls (0.74, 0.94, 0.98 for NA12878-specific calls) for L1HS, *Alu*Ya5/8, and *Alu*Yb8/9, respectively (Fig. 2, Table 1; Materials and Methods). A complete list of all annotated TE calls identified in NA12878 is available as a flat file on github (https://github.com/ekviky/TE-NGS).

**Fig.2.**
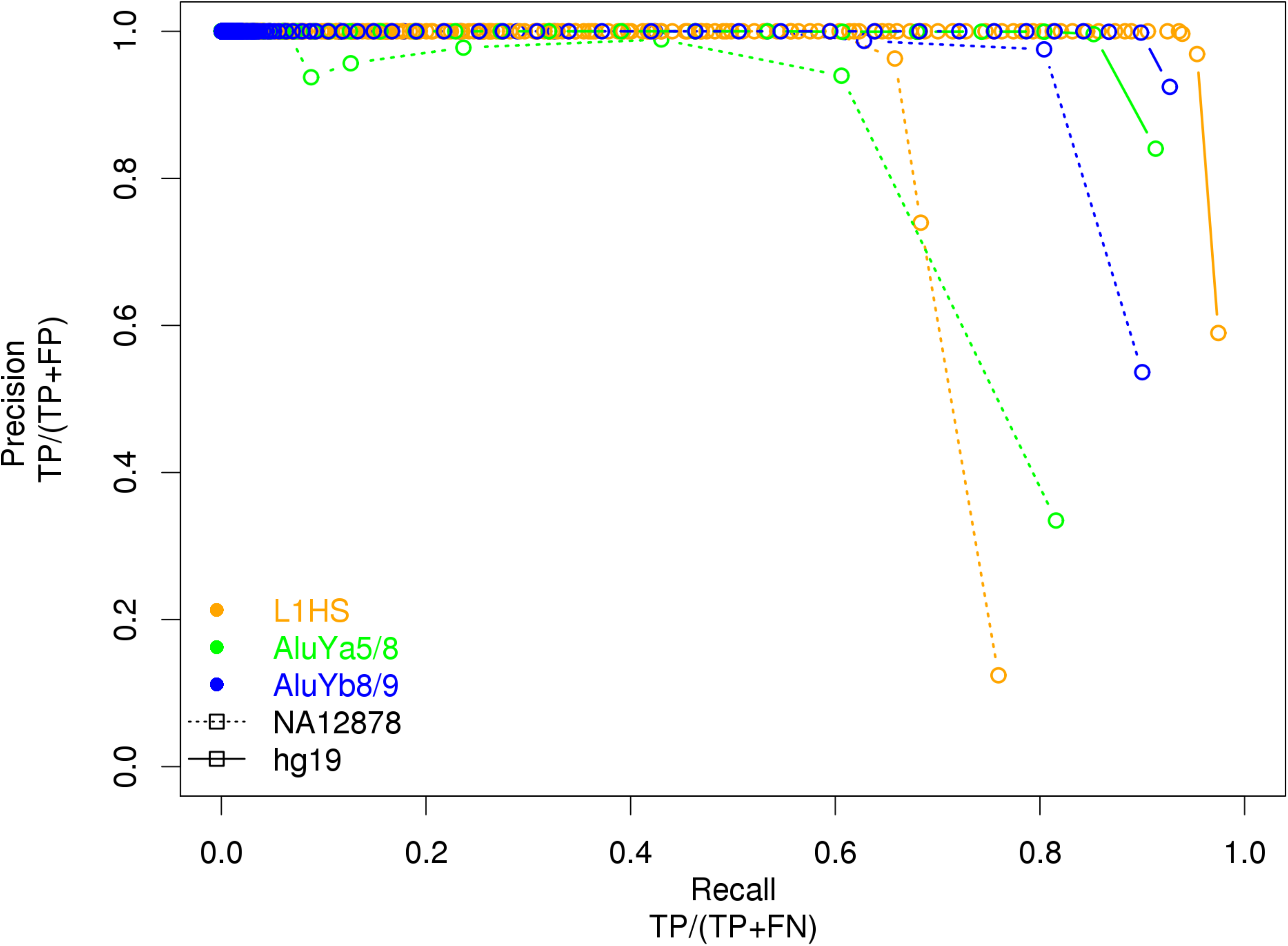
Precision and recall as a function of cluster read depth. Performance of TE insertion detection was assessed computing precision and recall separately for reference and polymorphic TEs present in NA12878. Reference (NA12878) true positives (TP) and false negatives (FN) were determined by comparison of clusters to reference (hg19) RepeatMasker annotations (NA12878-specific 1000 Genomes Phase 3 MEI calls), respectively. False positives (FP) were defined conservatively as TE candidate clusters failing to intersect any previously identified NA12878 non-reference events (see main text; Materials and Methods for details).

### Validation of NA12878 TE calls with long read data

We considered the possibility that a fraction of TE-NGS predictions in NA12878 that were classified as false positive could be caused by annotations missing from the 1000 Genomes Phase 3 MEI call set and the other call sets we considered. To address this possibility, we examined long-read sequence data for NA12878 for evidence of TE insertions at calls classified as FP. Long reads averaging >10,000 bp[34] are capable of spanning the entirety of a TE insertion in the case of *Alu*s (∼300 bp), and much if not all of LINE1 that are typically truncated upon insertion (∼500 bp and longer; [35]).

First, we obtained sequence data of NA12878 PacBio reads generated by the GiaB consortium [34, 36]. We interrogated these data for sequence signatures consistent with TE insertions. PacBio reads spanning FP calls and containing exact matches to TE-like sequences validated a total of 27/31 (0.87) FP as likely true insertions. By subfamily, 18/19, 7/9, and 2/3 L1HS, *Alu*Ya5/8, and *Alu*Yb8/9 FP calls, respectively, are represented by at least one read with TE-like sequence, with mean coverage depth of 31.26 (median 9) TE-like reads per locus.

Second, FP were manually inspected to detect evidence consistent with TE insertions using an orthogonal long read data set for NA12878 generated by Oxford Nanopore Technology (ONT, [37]). Visual inspection of these data confirmed that 21/31 (0.67) of FP calls are spanned by ONT reads consistent with TE insertion (Fig. S4, Materials and Methods).

Together, we find that at least one of the long-read data sets provided independent evidence supporting the presence of a TE insertion at 30/31 loci that were considered FPs in the initial analysis; 17/31 loci (10/19 *Alu*Ya5/8, 6/9 *Alu*Yb8/9 and 1/3 L1HS) were supported by evidence from both GiaB and ONT reads. This resulted in increased precision rates for both reference and polymorphic TE sets; using the more conservative estimate we find a precision of 0.985 (589/598), 0.999 (2335/2338) and 0.999 (1664/1666) for L1HS, *Alu*Ya5/8, *Alu*Yb8/9 reference calls respectively, and 0.86 (54/63), 0.98 (143/146), and 0.98 (119/121) for NA12878-MEI calls of the same types (Table 1). The 14 remaining FP calls represent a small fraction, 0.29% (14/4904), of total polymorphic TE predictions produced by this method.

### Application of TE-NGS to additional samples

Twelve additional clinical samples were sequenced by the TE-targeting method along with NA12878. These samples correspond to individuals with previously generated WES data and for whom no causal variants were identified. Table S4 summarizes the TE calls produced by application of TE-NGS. One individual (sample 8) failed all three targeting experiments and was excluded from further analysis. On average, in these samples we observe a fraction of 0.90, 0.87, and 0.88 of the expected reference L1HS, *Alu*Ya5/8, and *Alu*Yb8/9, respectively, similar to the observed rates of recall for NA12878 individual (see above).

We detected a total of 1,471 novel TE insertions, 1,438 of which are unique. By subfamily, this breaks down as an average of 42.5, 80, and 14 novel insertions of L1HS, *Alu*Ya5/8, and *Alu*Yb8/9 per individual (Table S4). Two percent (32/1438) of the novel events were observed in more than one individual, including two previously unreported insertions of L1HS that were observed in all 11 samples (chr2: 132998699-132998918; chr16: 61079102-61079356; both build GRCh37). The distribution of all identified novel insertions largely follows the genome-wide expectation, with no discernable chromosome bias nor enrichment near genes (data not shown).

#### Estimation of performance using trio data

Given the low estimated insertion rates (∼1 per 20 births for *Alu*; see above), we expect that the vast majority of novel TE insertions are rare polymorphic events inherited from either parent, and/or false positives, rather than *de novo* insertions. For two sets of trios included in this study, inheritance patterns were ascertained at TE loci observed in each proband, allowing us to estimate false positive and false negative calls. We evaluated sensitivity for the parental truth set (defined as clusters present in both parents), and novel TE insertions unique to the proband (absent in both parents) were considered as false positives calls for calculation of precision and recall (see Materials and Methods for details).

Of the 5,692 clusters present in both parents of Trio A, we observed 5,384 (95%) in the proband (sample 1); broken down by TE subfamily, the proposed method’s recall for this individual was 682/721 (0.95), 2972/3110 (0.96), and 1730/1861 (0.93) for L1HS, *Alu*Ya5/8, and *Alu*Yb8/9, respectively. For example, Fig. S5 illustrates a novel L1HS observed in all members of Trio A and thus inferred to be inherited in the proband. Of the total novel calls detected in this proband, 7/11 L1HS and 22/29 *Alu*Ya5/8 were absent from both parents and therefore classified as false positives, while every novel *Alu*Yb8/9 was inherited, corresponding to high rates of precision (>=0.99) for each assay (682/689, 2972/2994, 1730/1730 for L1HS, *Alu*Ya5/8, and *Alu*Yb8/9, respectively; Table S4; Materials and Methods).

The Trio B parental truth set contained 6,156 clusters, 5,756 (94%) of which were observed in the proband (sample 4; Table S4). In this particular individual, we observed 690/744 (0.93), 3258/3450 (0.94), and 1808/1962 (0.92) rates of recall for L1HS, *Alu*Ya5/8, and *Alu*Yb8/9, respectively. And similarly high rates of precision 0.89 (690/773), 0.94 (3258/3481), 0.98 (1808/1841) were obtained for L1HS, *Alu*Ya5/8, and *Alu*Yb8/9, respectively.

## DISCUSSION

We have developed a method for the identification of novel TE elements in human genomes, consisting of a molecular genomics protocol and bioinformatics pipeline (Fig. 1, S2). TE-NGS uses targeted NGS sequencing combined with genomic positional evidence to identify TE insertions, and novel loci are discriminated by further annotation with public databases. The approach here simultaneously targets the three elements responsible for the majority of active retrotransposition in the human lineage (*Alu*Ya5/8, *Alu*Yb8/9, L1HS). Using the well-characterized human sample NA12878 we show that the approach achieves high sensitivity and specificity (Fig. 2).

To determine the genomic locations of novel TE insertions, an assay must amplify not only the TE itself but the unique contextual sequence of each element. We achieved this by taking advantage of the Universal adapter primer present in Illumina genome libraries, using PCR to selectively amplify only the subpopulation of genomic molecules that span the junction of a TE insertion (Fig. 1).

The presence of many near-identical repeat inserts, and the large amount of genomic background sequences over TE-containing genomic molecules present challenges to generating on-target and usable data. In addition, molecules spanning inverted TE pairs in head-to-head orientation will propagate through the protocol, and are capable of hybridizing to Illumina flowcells, but as they consist of TE primer sites at both ends they contain only one of the two adaptors required for cluster formation, producing molecules ultimately incompatible with dual-adapter NGS sequencing and reducing the yield of the protocol. We used two incubations with an ssDNA exonuclease to systematically degrade these unwanted PCR products from the libraries (Fig. 1), allowing us to decrease genomic background DNA and remove molecules spanning inverted repeats. Introducing exonuclease digestion steps resulted in high proportions (87% or more; Table S3) of usable read data generated for each element.

The proposed protocol significantly improves upon on the total cost and sample usage compared to previous approaches. Current approaches require up to tens of millions of reads per sample *e.g.*, [21, 23, 24], offsetting the potential for deep multiplexing to decrease the burden of sequencing costs. By contrast, the protocol described here allowed us to sequence 39 libraries (13 samples x 3 elements) on a single Miseq lane. Given the levels of sensitivity attained (Fig. 2) and the complexity of the resulting libraries (averaging 143, 22 and 36 reads per L1HS, *Alu*Ya5/8, and *Alu*Yb8/9 locus, respectively; Table 1), we estimate that as many as 400 individuals could be multiplexed on a single lane of Illumina HiSeq 2500 (assuming a minimum of 200M reads, and ∼500,000 reads/sample). Moreover, only 10 ng of genomic library is required as input material; many library preparation kits perform well with as little as 5ng of genomic DNA (*e.g.,* New England Biolabs, Ipswich, Massachusetts).

Despite these gains, several areas remain for further improvement. The utility of the protocol would increase if the relatively high FN rate of ∼5% could be decreased. In part, FN are driven by sequence characteristics of the TE flanking sequences which influence the ability of the relatively short reads to map uniquely. Some FNs will be due to reference TEs that overlap CNVs in the particular individual. Unfortunately, Phase3 calls for TE deletions with respect to the reference genome are not readily available, such that a complete set of reference insertions known to be absent from NA12878 is not available at this time. Another cause of FNs is the variable coverage between individual TE loci, due to differential efficiencies in the various PCR steps in the protocol, which contribute to an underrepresentation of reads at some loci that could be mitigated by increasing the overall depth of sequencing.

We find that the observed detection rate for polymorphic NA12878 TEs is lower than that for Reference TEs. This could be due in part to the incomplete nature of Phase3 MEI annotations. For example, Phase3 MEI calls are annotated as LINE1 events, while our assay is designed to target specifically the active (Ta-0 and Ta subset) of L1HS subfamilies. It is possible that a proportion of these loci correspond to L1PA2 or other closely related subfamilies that are not enriched by TE-NGS (Table S3). Indeed, of the 35 full length LINE1 events in Phase 3 calls, we observe all but 2 (94%), consistent with recent assays targeting the “hot” and active subset of LINE1 events [25]. Furthermore, inspection of long-read data produced by both Pac Bio and ONT confirmed substantial proportions (18/19, 7/9, 2/3) of FP L1HS, *Alu*Ya5/8 and *Alu*Yb8/9 calls as likely true insertion events absent from the 1000 Genomes Phase3 call set.

The results of the PacBio and ONT long-read analysis also suggest that TE-NGS achieves high specificity for targeted TE subfamilies. Although the specificities presented here are conservatively calculated using calls validated by both PacBio and ONT reads, it is unlikely that positive evidence of a TE in a long and uniquely mappable ONT or PacBio read would occur by chance. Discrepancies between PacBio and ONT data could be explained by variable coverage in each data set, as well as a minority of unique insertions in the particular NA12878 cell lines utilized by each experiment.

The capacity to detect all TEs in an individual will require measures to sequence and align reliably in difficult genomic contexts. Although this protocol was developed for use with Illumina sequencing platforms, in theory, any existing NGS technology can be used insofar as the appropriate adaptor sequences and sequencing primers are publicly available for use in primer design. This has not been tested directly at this time, yet emerging long-read technologies may allow future adaptations of the protocol to sequence the TE and both its breakpoints in entirety. Long reads could also lead to enhanced sensitivities in difficult genomic regions (*e.g.*, nested insertions, inverted repeats).

Remaining challenges notwithstanding, the comprehensive nature of TE-NGS, its high sensitivity and specificity, and its cost-effectiveness due to the high fraction of on-target data, opens the possibility of adding effective TE screening to existing sequencing projects. This is particularly true for whole-exome sequencing projects, where TE-NGS can reuse part of the library preparation protocol to detect potential clinically relevant mutations that are not targeted by whole-exome methods. Importantly, the method described here provides an affordable assay for detecting novel TE insertions, a significant source of structural variation in human genomes that is not currently investigated in detail and may be more relevant in clinical settings than is currently appreciated.

## MATERIALS AND METHODS

### Samples

We obtained NGS libraries previously sequenced by the Oxford Genomics Centre from samples recruited as part of the Oxford Biomedical Research Centre’s Clinical Exomes project; the NA12878 library was kindly provided by Oxford Genomics Centre.

### TE Primer Design

To selectively enrich NGS libraries for TE-containing fragments, PCR primers were designed to amplify actively transposing members of *Alu* (Yb8/9 and Ya5/8) and L1HS (Ta-1 or 0) subfamilies (Table S2). Primers were designed using Primer3 software with the following characteristics: (i) targeting diagnostic nucleotides specific to each subfamily; (ii) positioned in proximity to 3’ tail of the consensus sequence to optimize coverage of flanking sequence; and (iii) containing low complementarity to facilitate multiplexing in single PCR reactions [38]. Each element was targeted by two primers. *First*, two TE-target primers were designed to amplify the subfamilies of interest, one primer each for *Alu* and LINE1. *Second*, three TE-nested primers complimentary to subfamily-specific bases located further downstream (3’) from the targeting primer. The L1HS-target primer, L1HsTailSp2, was previously published [21], Table S2).

Primer subfamily specificity was measured by determining the number and locations of primer matches in the reference genome (hg19) allowing at most 1 mismatch at positions excluding the 3’ ultimate nucleotide, and intersecting matches with RepeatMasker annotations [29] obtained from UCSC Table Browser [39].

TEs located in head-to-head orientation, i.e. inverted repeats capable of exponential amplification, were identified by defining genomic positions of all possible combinations of TE-target primer and a primer’s reverse complement (likewise TE-nested primer and reverse complement). Using the same defined primer match parameters (see above) and limiting primer distances to within a maximum of 1-kb (estimated maximum library fragment size), we identified 352 loci as inverted repeats.

### TE-NGS Library Preparation

#### Preparation of genomic libraries

First, genomic molecules were sheared to an average library insert size of ∼300 bp using Covaris following manufacturers specifications (Covaris, Woburn, Massachussets). Library construction consisting of end repair, adapter ligation and PCR extension was performed using NEBNext Ultra DNA Library Kit for Illumina (New England Biolabs, Ipswich, Massachusetts; cat. E7370) to generate libraries with full-length adapters including index 1 (i7) for Illumina paired-end sequencing.

#### TE-target amplification

TE-containing fragments were exponentially amplified from genomic library using the two TE-targeting primers in conjunction with the Illumina Universal PCR primer (P5) in a multiplex reaction (Table S2). The PCR reaction was performed in a 50 uL volume containing NEB Q5 High Fidelity 2x PCR mastermix (New England Biolabs; cat. M0541L), each TE primer at concentration of 0.5uM, 1uM of Illumina Universal P5 PCR primer, 2mM of magnesium, and ∼10 ng of library DNA. PCR cycling conditions were as follows: initial denaturation at 98C for 2 minutes; 10 cycles of (denaturation at 98C for 10 sec, annealing at 67C for 30 sec, extension at 72C for 30 sec); and final extension at 72C for 5 minutes.

#### Post-PCR DNA Purification

DNA purification and library size selection was performed by solid phase reversible immobilization (SPRI) using Agencourt AMPure XP magnetic beads (Beckman Coulter, Brea, California; cat. A63882). A ratio of 1.8x beads to reaction volume was used to selectively retain fragments ∼100 bp and larger, and to ensure removal of Illumina P5 primers from the reaction solution.

#### TE-target asymmetric amplification & digestion of background ssDNA fragments

To asymmetrically amplify fragments containing TE-targeting primer sites, a PCR reaction was performed with NEB Q5 High Fidelity 2x PCR mastermix, both TE-target primers at 0.5 uM each, 2mM magnesium, and clean eluted PCR reaction from previous step. Cycling parameters were denaturation at 98C for 2 min, followed by 2 cycles of (denaturation at 98C for 10 sec, annealing at 67C 30 sec, and extension at 72C 30 sec).

Double-stranded DNA (dsDNA) products at this step include linear amplification from TE targets, and exponential amplification of TE targets oriented in head-to-head fashion. Background genomic DNA molecules lacking TE-target priming sites remain largely denatured (due to their high complexity), in single-stranded DNA (ssDNA) form. To remove the unwanted ssDNA fragments from the library after asymmetric amplification, libraries were incubated with RecJf exonuclease (New England Biolabs, cat. M0264L) following manufacturer’s specifications.

#### Post-digest DNA Purification

Libraries following exonuclease digestions were purified by SPRI using AMPure XP magnetic beads in a 1.8x ratio of beads to reaction volume, to selectively retain fragments ∼100 bp and larger and to remove TE-target primers from the reaction solution.

#### Illumina adapter (P5) asymmetric amplification & digestion of head-to-head ssDNA fragments

To asymmetrically amplify fragments containing the Illumina universal adaptor sequence, a linear PCR reaction was performed with Illumina P5 PCR primer at 0.5 uM, NEB Q5 High Fidelity 2x PCR mastermix, and clean eluted PCR reaction from previous step. Cycling parameters were denaturation at 98C for 2 min, followed by 2 cycles of (denaturation at 98C for 10 sec, annealing at 63C 30 sec, and extension at 72C 30 sec).

At this step, dsDNA products will result from linear amplification of TE targets. Molecules containing TEs in head-to-head orientation lack Illumina universal adapters, and remain largely denatured in ssDNA form, for subsequent removal by exonuclease digestion. Digestion was performed by incubation with RecJf exonuclease as above.

#### TE-nested amplification

To further increase specificity, three individual PCR reactions were performed each targeting a specific subfamily. TE-nested primers were designed for further TE specificity, and contain nucleotides to partially reintroduce the Illumina index i7 adapter that contains a unique 6-mer index per TE, per sample (Table S2). Each PCR reaction was performed in a separate 50 uL volume containing a 4ul aliquot of the post-digest library from previous step, NEB Q5 High Fidelity 2x PCR mastermix, one TE-nested primer at concentration of 0.5uM, and Illumina P5 Universal adapter primer at 0.5uM. PCR cycling conditions were as follows: initial denaturation at 98C for 2 minutes; 15 cycles of (denaturation at 98C for 10 sec, annealing at Ta_primer for 30 sec, extension at 72C for 30 sec); and final extension at 72C for 5 minutes; where Ta_primer corresponds to (68C L1HS-nested, 68C *Alu*Ya58-nested, and 64C *Alu*Yb89-nested primer, respectively; see Supplemental Material for detailed procedures; Table S1).

#### Post-PCR DNA Purification

Libraries following nested amplification were purified by SPRI using AMPure XP magnetic beads in a 1.8x ratio of beads to reaction volume, to selectively retain fragments ∼100 bp and larger.

#### Adapter extension

Enrichment for molecules containing full-length double-stranded DNA adapter libraries was achieved by PCR adapter extension. A 50uL PCR reaction was performed with Illumina P5 Universal PCR primer and P7 index primers at 0.5 uM each, NEB Q5 High Fidelity 2x PCR mastermix, and clean eluted PCR reaction from previous step. Cycling parameters were denaturation at 98C for 2 min, 5 cycles of (denaturation at 98C for 10 sec, annealing at 65C for 30 sec, and extension at 72C for 30 sec), and final extension at 65C for 5 minutes.

#### Library multiplexing and sequencing

The resulting TE-enriched libraries were purified, quantified then pooled according to relative molarities. Full length adapter libraries were purified by SPRI using AMPure XP magnetic beads in a 1x ratio of beads to reaction volume, to selectively retain fragments ∼150 bp and larger. The molarity of each library was computed by obtaining the concentration using Qubit (Life Technologies, Carlsbad, California) and size distribution on Agilent Tapestation High Sensivity D1000 Screen Tape (Agilent Technologies, Santa Clara, California; cat. 5067-5584). Fig. S1 demonstrates the fragment distribution of a typical library, that is free of short (<100 bp) fragments and displays a peak at ∼200 bp, or approximately half the size of the starting library.

TE-enriched libraries were subsequently multiplexed according to relative molarities (see Supplemental Material). The final library pool was diluted to 10 nM and sequenced by 151-bp paired-end reads using Illumina MiSeq (Illumina Inc, San Diego CA).

### Identification of TE loci from Targeted Sequencing

#### Alignment

Sequence data generated by TE-enrichment were de-multiplexed and aligned to the human reference genome (hg19) using the Burrows-Wheeler Aligner [40] followed by refinement of BWA alignments with Stampy [41], sorting of BAM alignments using Samtools sort [42], removal of PCR duplicates using the module MarkDuplicates in Picard [43] and BAM indexing using Samtools index.

#### Post-alignment preprocessing of reads

TE loci in the targeted sequencing data were identified by a bioinformatic pipeline that incorporates signatures of TE loci such as (i) primer matches (ii) matches to “TE-like” sequence (iii) read orientation (Fig. S2). Amplification of TEs by TE-NGS is performed with the TE-nested primer in conjunction with the Universal adapter primer (P5), that is used to produce Illumina’s “read 1”. Therefore, Illumina read 1 (derived from Illumina i5/P5 primer) will be generated from the unique flanking sequence, while Ilumina “read 2” (produced from i7/P7 primer) will contain primer, 3’ portion of TE, and poly-A tail sequences (Fig. 1). As such, different requirements were placed on the two reads generated from the targeted paired-end sequencing as follows. Reads derived from the mobilome were distinguished from genomic background by requiring exact matches to the full-length TE-nested primer sequences at the start of read 2. This strict filtering minimized the potential for false positives calls. Read pairs with read 2 lacking any discernable match to the primer (defined as exact matches to first 7/10 nucleotides of primer) correspond to genomic background and were removed from further analysis.

So-called TE-like sequences were defined as sequence downstream from the TE-nested primer, *i.e.* (3’) relative to the TE sequence, and excluding the poly-A tail. TE-like sequence was determined for each TE subfamily by analysis of consensus sequences obtained from Repbase [30]. Multiple alignments were produced separately for all *Alu* and all LINE1 subfamily sequences using MUSCLE [44]. Based on the position of the TE-nested primer in the consensus sequence, the length of TE-like sequence varies for each subfamily (8, 37, and 29 nucleotides for L1HS, *Alu*Ya5/8 and *Alu*Yb8/9, respectively). Read 2 reads with a maximum number of allowed mismatches to the TE-like sequence (3, 10, and 10 for L1HS, *Alu*Ya5/8 and *Alu*Yb8/9, respectively) were considered TE-derived and their read pairs retained (Fig. S6).

TE-enriched libraries are anticipated to include sequences generated from TE insertions absent from the reference genome. Read 2 sequences generated from short library fragments could potentially contain TE-nested primer sites, 3 prime TE and poly-A tail sequences either failing to map and/or capable of mapping to near-identical sequences contained in TEs present in the reference at alternative locations, including potentially on different chromosomes. Therefore, unlike standard genomic library processing, we placed no requirements for alignments to adhere to Sam specification’s flags for both mates mapping, nor mapping as “proper pair”, and placed no restrictions on the insert size. Unmapped reads and reads with mapping quality 0 were removed after primer and TE-like sequence requirements, to ensure that TE loci spanned by clusters of read 1 mapping reliably (minimum MAPQ>=3) were retained for analysis.

#### Clustering

To identify unique TE insertions we created a catalog of filtered reads covering a genomic coordinate interval (cluster), and second, used mapping signals in conjunction with genome annotations to classify predicted insertion calls.

In the first step, individual reads were clustered by genomic position for each library of each uniquely indexed sample and TE primer combination. Clusters were annotated with various attributes of the corresponding catalog of reads for use as evidence at each locus in downstream filtering. Clusters were generated requiring a minimum inter-distance of 200 bp between neighboring clusters.

Clusters with a minimum cluster size based on sequence read length (100 bp) were retained, and clusters were required to have a minimum of 2 reads derived from the TE’s unique flanking sequence (read 1), having minimum mapping quality (MAPQ>=3).

Clusters in regions of potential noise due to read mapping artifacts were filtered. Calls in proximity to annotated reference gaps (unassembled poly-Ns; 500-bp window), regions enriched for satellite, i.e. centromeres, and sub-telomeres (defined as 1-Mb window), and located on chrY were excluded from downstream analysis (annotations obtained from USCS Table Browser; [39]).

#### Annotation

In the second step, cluster loci were annotated using known TE databases [29, 31] and insertions from previously published TE assays [14, 15, 21, 22]. To achieve this, we compiled a comprehensive local database of polymorphic *Alu* and LINE elements called polyTEdb. The compiled polyTEdb is available for download from github (https://github.com/ekviky/TE-NGS). Clusters were intersected with annotations and labeled with TE annotation subfamily (when available); clusters mapping to within 600 bp window of the TE 3’ end (when strand orientation provided) were annotated.

Calls produced by the above pipeline were classified as *reference* if reference genome contains insertion at the position and matches subfamily of target assay; *known non-reference* if evidence of an insertion of polyTE at that position in polyTEdb (because subfamily-specific information is not routinely/consistently provided). All calls lacking previous evidence of a TE insertion at that position were considered *novel*.

### Reference TE in NA12878

Targeted loci present in hg19 were defined by identifying all matches to TE-target primers, and TE-nested primers using the parameters defined above (see Primer Design). TEs containing matches to both primers, within a maximum distance of 200 bp, and in proper order and orientation with respect to reference strand, were classified as targeted loci. Targeted loci excluding chrY, annotated reference gaps (unassembled poly-Ns; 500-bp) or centromeres/sub-telomeric regions (defined as 1-Mb window) were retained in the Reference TE set (annotations obtained from UCSC Table Browser; [39]). Loci were further refined to exclude target loci overlapping NA12878-specific copy number variants (CNV) as annotated in Database Genomic Variants [45]. This resulted in 624, 2,739, and 1,847 reference L1HS, *Alu*Ya5/8, and *Alu*Yb8/9 insertions respectively (Table 1).

### NA12878-specific TE

Mobile element insertions (“MEI”) observed in the NA12878 individual were parsed from variant vcf files obtained from Phase3 of 1000 genomes [14, 46]. 857 *Alu* and 76 LINE1 remained after filtering to exclude annotated reference gaps, centromeres/sub-telomeric regions, or NA12878-specific CNV (see above definitions). Of the 857 *Alu* loci, 163 *Alu*Ya5/8 and 61 *Alu*Yb8/9 included subfamily designation; these insertions were used to define respective *Alu* false negatives in the TE-NGS call set; all 76 LINE1 calls lacked subfamily information and were used to define L1HS false negatives.

### Parental Truth Sets

For two trios included in this study, inheritance patterns were ascertained for TEs observed in each proband as follows. The parental truth set for each trio was defined as TE insertions detected at the same location (to within a window of 100 bp) in both parents; 5,692 TE were present in both parents of Trio A, and 6,156 for Trio B.

False positives in the proband of each trio were classified as novel TE insertions absent from both parents (i.e., apparent “de novo”). Because we observe virtually every TE in a genome with a minimum of one read (Fig. 2; see above), in defining false positives we relaxed the minimum read coverage requirement for the parental clusters and conservatively defined *de novo* calls in the proband as novel loci lacking evidence from even one read in one parent.

### Precision and Recall

To characterize the ability of TE-NGS to predict true insertions, we evaluated the sensitivity or recall, computed as TP/(TP + FN), for each of the above truth sets. Precision, computed as TP/(TP + FP), was used to characterize the ability of the assay to reject false insertion predictions.

### Validation of NA12878 calls with Long Read Data

Loci corresponding to potential false positive calls were inspected with two data sets generated by long-read sequencing platforms. BAM alignments provided by Genome in a Bottle (GiaB) using PacBio sequencing data were obtained from [47]. Long reads spanning TE calls were required to contain exact matches to TE-like sequences (see above).

BAM alignments of NA12878 reads generated by ONT were obtained from the Wellcome Trust Centre for Human Genetics [37]. Alignments were manually inspected by Integrated Genome Viewer (IGV;[48]. FP loci were inspected for characteristics typical of TE insertions: stretches of inserted sequence and/or clipping of long reads containing sequence absent from the reference (see Fig. S4).

## DECLARATIONS

### Ethics approval and consent to participate

Individual researchers had explicit research consent to undertake genetic investigation into the cause of the relevant disease. Samples provided by Anna Schuh were approved by Oxford Research Ethics Committee C (REC reference: 09/H0606/5); samples provided by Ed Blair were approved by NRES West Midlands, Coventry and Warwickshire (REC reference 13/WM/0466). The samples provided by Andrew Wilkie were approved by London Riverside Research Ethics Committee (REC reference 09/H0706/20). Informed consent for participation in this study was obtained from healthy donors, patients, or their parents in all cases. Further information and documentation to support will be made available to the Editor on request.

### Consent for publication

Not applicable

### Availability of data and material

The datasets generated and analysed during the current study are available in the TE-NGS repository, hosted on (https://github.com/ekviky/TE-NGS).

### Authors’ contributions

EK, GL, PP designed the TE targeting protocol; EK, GL designed the TE detection algorithm. EK conducted the experiments, designed and wrote the code implementing the algorithm. EK, GL analyzed, interpreted the results, and wrote the manuscript. JCT provided samples. EK, PP, JCT, GL wrote, reviewed, and edited the manuscript. All authors read and approved the final manuscript.

## Acknowledgements

We thank the patients and their families who consented to these studies. We are grateful to Andrew Wilkie, Anna Schuh, and Ed Blair for permission and use of samples. Sequencing data was generated by the High-Throughput Genomics Group at the Wellcome Trust Centre for Human Genetics. We would like to thank HTG and the Library Prep team in particular for generous use of lab space and helpful discussions. We thank Amy Trebes, Samantha Knight, Kateryna Makova, and XX anonymous reviewers for helpful comments on an earlier version of this manuscript. The views expressed in this manuscript are those of the authors and not necessarily the Wellcome Trust and Department of Health.

